# A genetic strategy to measure insulin signaling regulation and physiology in *Drosophila*

**DOI:** 10.1101/2022.05.31.494190

**Authors:** Deborah D. Tsao, Kathleen R. Chang, Lutz Kockel, Sangbin Park, Seung K. Kim

**Affiliations:** Department of Developmental Biology, Stanford University School of Medicine, Stanford, CA; Department of Medicine (Division of Endocrinology, Metabolism, Gerontology), Stanford University School of Medicine, Stanford, CA; Department of Pediatrics (Division of Endocrinology), Stanford University School of Medicine, Stanford, CA; Stanford Diabetes Research Center, Stanford University School of Medicine, Stanford, CA

## Abstract

Insulin regulation is a hallmark of health, and impaired insulin signaling promotes metabolic diseases like diabetes mellitus. However, current assays for measuring insulin signaling in all animals remain semi-quantitative and lack the sensitivity, tissue-specificity or temporal resolution needed to quantify *in vivo* physiological signaling dynamics. Insulin signal transduction is remarkably conserved across metazoans, including insulin-dependent phosphorylation and regulation of Akt/Protein kinase B. Here, we generated transgenic fruit flies permitting tissue-specific expression of an immunoepitope-labelled Akt (AktHF). We developed enzyme-linked immunosorption assays (ELISA) to quantify picomolar levels of phosphorylated (pAktHF) and total AktHF in single flies, revealing dynamic tissue-specific physiological regulation of pAktHF in response to fasting and re-feeding, exogenous insulin, or targeted genetic suppression of established insulin signaling regulators. Genetic screening revealed *Pp1-87B* as an unrecognized regulator of Akt and insulin signaling. Tools and concepts here provide opportunities to discover tissue-specific regulators of *in vivo* insulin signaling responses.

**Author Summary:** Insulin is an essential hormone that controls metabolism in all animals, by regulating energy use and growth of target tissues. Impaired insulin signaling (“resistance”) in humans underlies development of type 2 diabetes, a pandemic disease causing significant morbidity and mortality. The genetic risk for insulin resistance is complex, and studies of diabetes are limited by a lack of tools to measure insulin resistance in a sensitive, quantitative, and tissue-specific way. Here, we describe a new technique to measure the strength of insulin signaling in fruit flies. By combining fruit fly genetics with antibody-based assays, we can quantify phosphorylated Akt, an evolutionarily conserved target of insulin signaling, in specific tissues of the adult fly. We show this technique can detect changes in insulin signaling after fasting and refeeding, addition of exogenous insulin, or genetic disruption of the insulin signaling pathway. We used this method to discover a new regulator of insulin signaling, a phosphatase enzyme encoded by *Pp1-87B*. This exciting new tool should advance our ability to study and discover additional regulators of insulin signaling and resistance.

## Introduction

Insulin is a crucial regulator of metabolism in multicellular animals, including the fruit fly *Drosophila melanogaster* [1]. Altered insulin sensitivity – like insulin resistance in liver, adipose and other insulin target tissues – can lead to pancreatic β cell failure and promote the pathogenesis of type 2 diabetes mellitus (T2DM) in humans [2-4]. Studies in patients with obesity and/or T2DM convincingly demonstrate a genetic basis for the pathogenesis of declining insulin sensitivity [5]. However, investigations of T2DM genetics are challenged to link altered gene function to *in vivo* phenotypes, and identify tissue(s) where gene function is required. *Drosophila* provides unique opportunities to dissect insulin signaling and resistance by combining genetics with appropriate physiological assays, like quantification of insulin signaling in target tissues. Moreover, the evolutionary conservation of this pathway could lead to discovery of novel insulin signaling regulators in mammals and humans.

*Drosophila* research has made important contributions to elucidating molecular and genetic regulation of insulin signaling in larval and adult organs [6, 7], but progress in this area has also been hampered by a reliance on qualitative or semi-quantitative assays [8-10]. For example, signaling through insulin receptor (IR) and IR substrates (IRS1/2) leads directly to phosphorylation of Akt/Protein kinase B (hereafter called Akt), but assessment of this conserved feature of insulin signaling is limited in *Drosophila* to immune-staining of phospho-Akt (pAkt) in preserved tissues, or western blotting [9, 10]. Similar limitations hamper studies of insulin signaling in other experimental systems, including mammals. An ideal *in vivo* assay for pAkt or Akt would permit absolute quantification from specific tissues at picomolar levels, but to date there is no such assay for flies or other metazoans.

Here, we describe generation of unique fly strains expressing *Drosophila* Akt tagged with two immuno-epitopes (Hemagglutinin and FLAG: hereafter, AktHF). We developed ELISA methods to quantify phospho-AktHF (pAktHF) and total AktHF. Directed expression of a transgene encoding AktHF facilitated tissue-specific measures of pAktHF and total AktHF from single flies. These measures provided unprecedented *in vivo* assessments of insulin signaling and regulation by genetic and physiological mechanisms, and led to identification of previously undetected regulators of insulin-dependent Akt signaling. Our study provides potent new tools for studying insulin signaling, physiology and mechanisms underlying development of insulin resistance, and diseases like diabetes mellitus.

## Results

### A novel ELISA to measure *in vivo* Akt phosphorylation

To enable quantification of phospho-Akt (pAkt) in flies, we developed an enzyme-linked immunosorbent assay (ELISA) for an immune-epitope tagged *Drosophila* Akt protein (**Figure 1A**) that can be expressed in any tissue/organ of interest using a tissue-specific binary expression system (either LexA/LexAop or Gal4/UAS; Methods). First, we generated and linked a transgene encoding Akt incorporating HemagglutininA (HA) and FLAG at the carboxy terminus (**Figure 1A**) to the LexA-responsive upstream activating sequence (*LexAop-AktHF*), and introduced a single copy of this construct into the *Drosophila* genome by site-directed insertion (Methods). To prevent unwanted effects of Akt misexpression in developing tissues, we used the temperature-sensitive Gal80 (*Gal80*^*TS*^) to restrict expression of AktHF to adult flies. In adult *Drosophila*, the fat body is an essential organ governing energy homeostasis that can store circulating fatty acids and glucose in response to insulin stimulation; thus, we focused our initial studies on the phosphorylation of Akt in this tissue. To this end, we generated a triple-transgenic fly harboring *r5-LexA, Tubulin-Gal80*^*TS*^, and *LexAop-AktHF* where AktHF expression in adult fat body can be modulated by a temperature shift from a restrictive 18°C to a permissive 30°C. We observed that these flies have normal growth, developmental timing, and fertility compared to control single transgenic *LexAop-AktHF* flies or *r5-LexA* flies maintained at 18°C.

**Figure 1:**
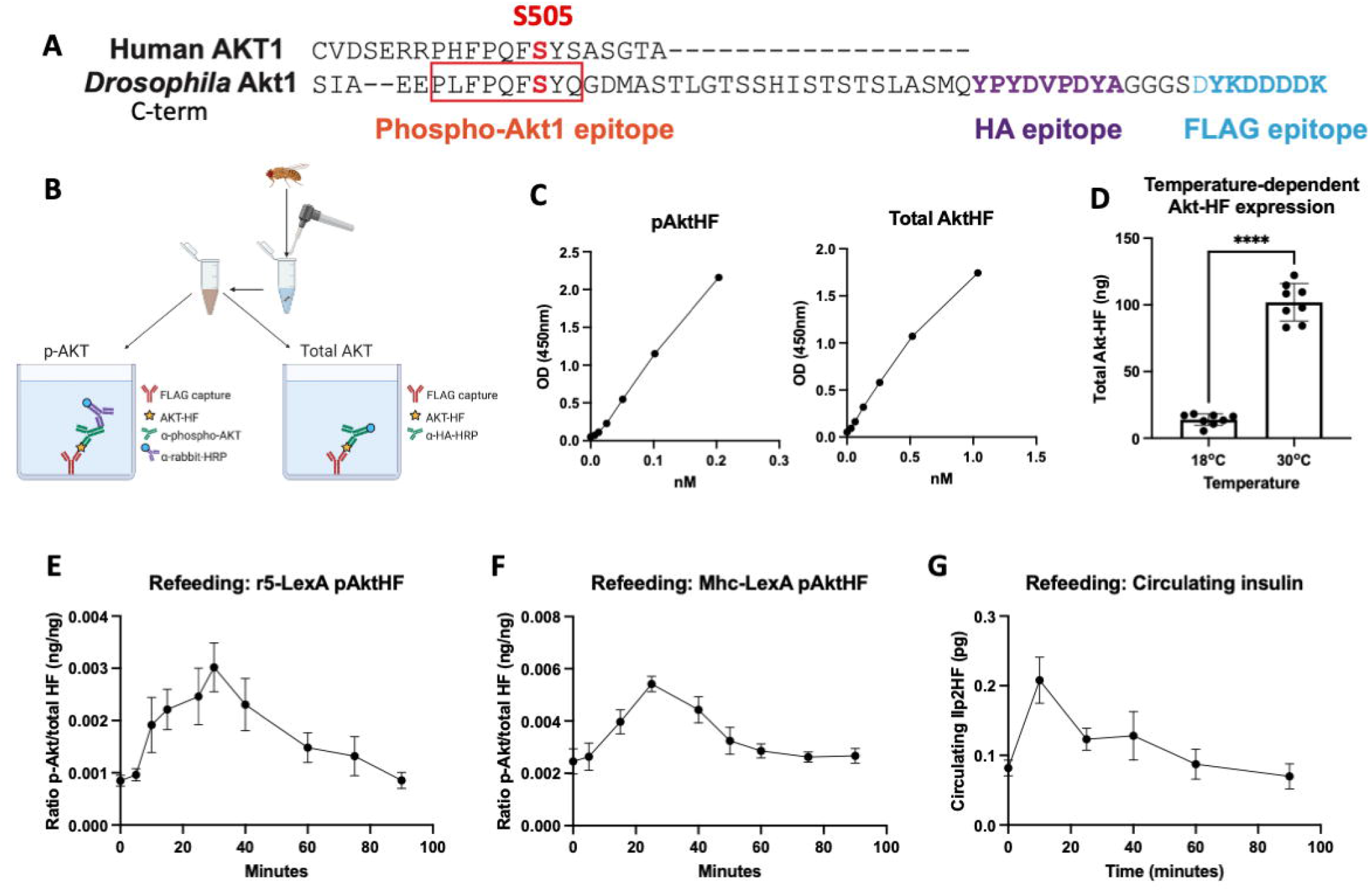
*Drosophila* pAktHF ELISA A) HA- and FLAG-tagged *Drosophila* Akt1 (AktHF), showing the conserved serine 505 phosphorylation site. B) Schematic summary of the two parallel sandwich ELISAs for phosphorylated and total AktHF in a single *in vivo* adult fly. C) Concentrations of phosphorylated and total AktHF were quantified by ELISA using phospho-Akt standard peptide (PLFPQFpSYQGDYKDDDDK, 0 to 0.5ng/mL, 0.2035nM) or FLAG(GS)HA standard peptide (DYKDDDDKGGGGSYPYDVPDYA, 0 to 2.5ng/mL, 1.036nM). D) The expression of AktHF is under control of Gal80^TS^, shown at the inhibitory 18°C and the permissive 30°C (n=8 flies per condition), two-tailed *t*-test *P*<0.0001. E) Oral glucose-stimulated induction of AktHF phosphorylation in adult fat body (n=4 flies per time point). F) Oral glucose-stimulated induction of AktHF phosphorylation in the adult muscle (n=4 flies per time point). G) Oral glucose-stimulated insulin secretion and clearance in adult flies (n=6 flies per sample, 3 samples per time point).

To evaluate the expression level of AktHF in adult fat body, we homogenized individual triple-transgenic flies, then conducted two parallel ELISAs to measure total AktHF and phospho-AktHF (pAktHF) in lysates. For both ELISAs, we used an anti-FLAG capture antibody (Methods: **Figure 1**). Total AktHF was measured by ELISA using an anti-HA antibody, while pAktHF was measured using an anti-pAkt antibody recognizing phosphorylated serine 505 of *Drosophila* Akt (**Figure 1A, B**); this antibody also recognizes the orthologous phosphoserine 473 of Akt1 in mice and humans (Methods). To quantify the signal detected in these parallel assays, we synthesized two peptide ‘standards’: FLAG(GS)HA peptide for total AktHF quantification, and PLFPQFpSYSA(GS)FLAG peptide for phosphorylated AktHF quantification (Methods). Both total and pAktHF were measured in a linear range from 5-10 pM or higher (**Figure 1C**). Unphosphorylated Akt505 peptide standards were not detected in the phospho-Akt ELISA (Methods), demonstrating the specificity of this ELISA for phospho-AktHF and total AktHF quantification. We compared flies reared and maintained at 18°C, or reared at 18°C and maintained at 30°C for 48 hours, and observed induction of AktHF expression at 30°C (**Figure 1D**). Thus, conditional expression of AktHF in adult fat body permitted quantification of total AktHF and pAktHF in a single fly at picomolar levels.

### Dynamic physiological regulation of Akt phosphorylation measured by ELISA

Pancreatic insulin secretion evoked by a meal stimulates Akt phosphorylation in mammalian target organs like liver, adipose and muscle [1]; thus, to assess the physiological relevance of our Akt/pAkt reporter system, we measured nutrient-dependent and insulin-dependent induction of pAkt. Adult flies expressing *LexAop-AktHF* specifically in fat body were subjected to a fasting and glucose refeeding challenge. After glucose re-feeding, pAkt levels rose for 30 minutes then declined thereafter to fasting levels by 90 minutes (**Figure 1E**). We observed similar dynamics in flies expressing *LexAop*-*AktHF* in muscle (**Figure 1F**), using the adult muscle-specific Mhc-LexA driver. Although the timecourse and magnitude of pAkt changes were comparable between tissues, the ratio of phosphorylated-to-total AktHF was greater in muscle than fat body. In these studies, total Akt did not significantly change during the timecourse tested (**Figure S1**), and we normalized pAkt to total Akt levels. Measures of circulating insulin levels in flies expressing Ilp2HF [11] showed that the shape and duration of pAkt induction corresponded well with the dynamic increase then clearance of circulating Ilp2HF in this fasting-refeeding paradigm (**Figure 1G**). Thus, we can detect dynamic, nutrient-dependent regulation of *in vivo* pAkt levels in crucial organs regulating metabolism on a timescale of seconds to minutes.

To assess the dependence of Akt phosphorylation on insulin signaling, we developed an analogous *ex vivo* pAktHF/AktHF ELISA assay based on addition of purified insulin (**Figure 2A**; Methods). After insulin addition to dissected abdominal fat body from individual *r5-LexA/LexAop-AktHF* flies, we observed rapid increase, then plateau of the pAktHF/total AktHF ratio within 10 minutes (**Figure 2B**). As expected, the amount of exogenous insulin determined the peak levels pAktHF in a dose-dependent manner (**Figure 2C)**. These findings provide strong evidence for insulin-dependent phosphorylation of AktHF and, together with our *in vivo* findings, support the view that the AktHF ELISA provides rapid, reproducible measures of insulin signaling and Akt phosphorylation in single adult flies, with unprecedented tissue specificity on a physiologically relevant timescale.

**Figure 2:**
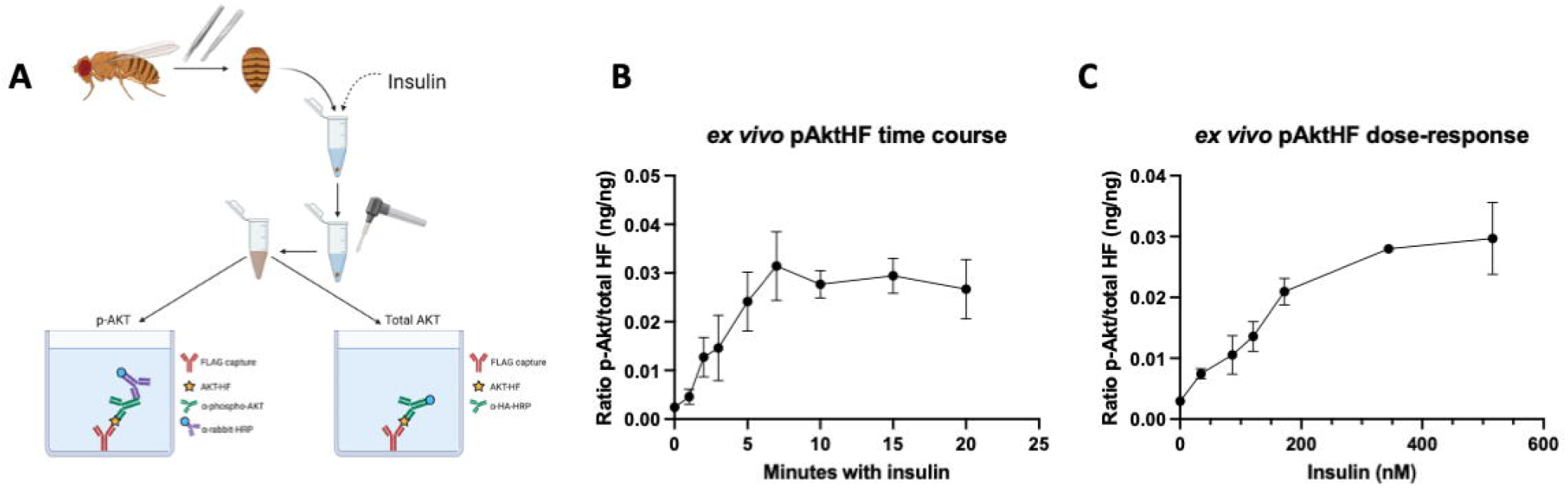
pAktHF *ex vivo* assay A) Schematic summary of the *ex vivo* AktHF ELISA, showing the dissection of the adult fat body and the addition of exogenous insulin. B) Insulin-stimulated phosphorylation of AktHF over time. (172nM human insulin, n=4 fat bodies per time point). C) Dose-response curve for human insulin, assessed by the phosphorylation of AktHF (n=4 fat bodies per concentration).

### Measuring dynamic changes of insulin signaling and sensitivity by pAktHF ELISA

Insulin signaling in target organs is altered by changes in insulin output, but these physiological signaling dynamics have not been previously measured in adult flies [1]. Here, we modulated insulin output in insulin producing cells (IPCs), then measured changes in pAktHF in the fat body. To simultaneously express transgenes in both IPCs and the fat body, we combined the Gal4/UAS and LexA/LexAop systems (**Figure 3A-H**). With *Ilp2-Gal4* driving expression of UAS-transgenes in the IPCs and *r5-LexA* driving expression of *LexAop-AktHF* in the fat body, we could simultaneously perturb IPC function and measure effects on fat body insulin signaling (**Figure 3E**). As previously reported [11], conditional expression of the inward-rectifying potassium channel KCNJ2 (Kir2.1) in IPCs (Methods, **Figure 3A**) led to durable reduction of circulating Ilp2HF (**Figure 3A-D**). Within one day of KCNJ2 expression, we observed reduced *in vivo* pAktHF levels in fat body (**Figure 3F**). However, this effect waned over time and after 9 days of IPC silencing by KCNJ2, we found that *in vivo* pAktHF levels were indistinguishable from controls with normal insulin levels (**Figure 3G,H**).

**Figure 3:**
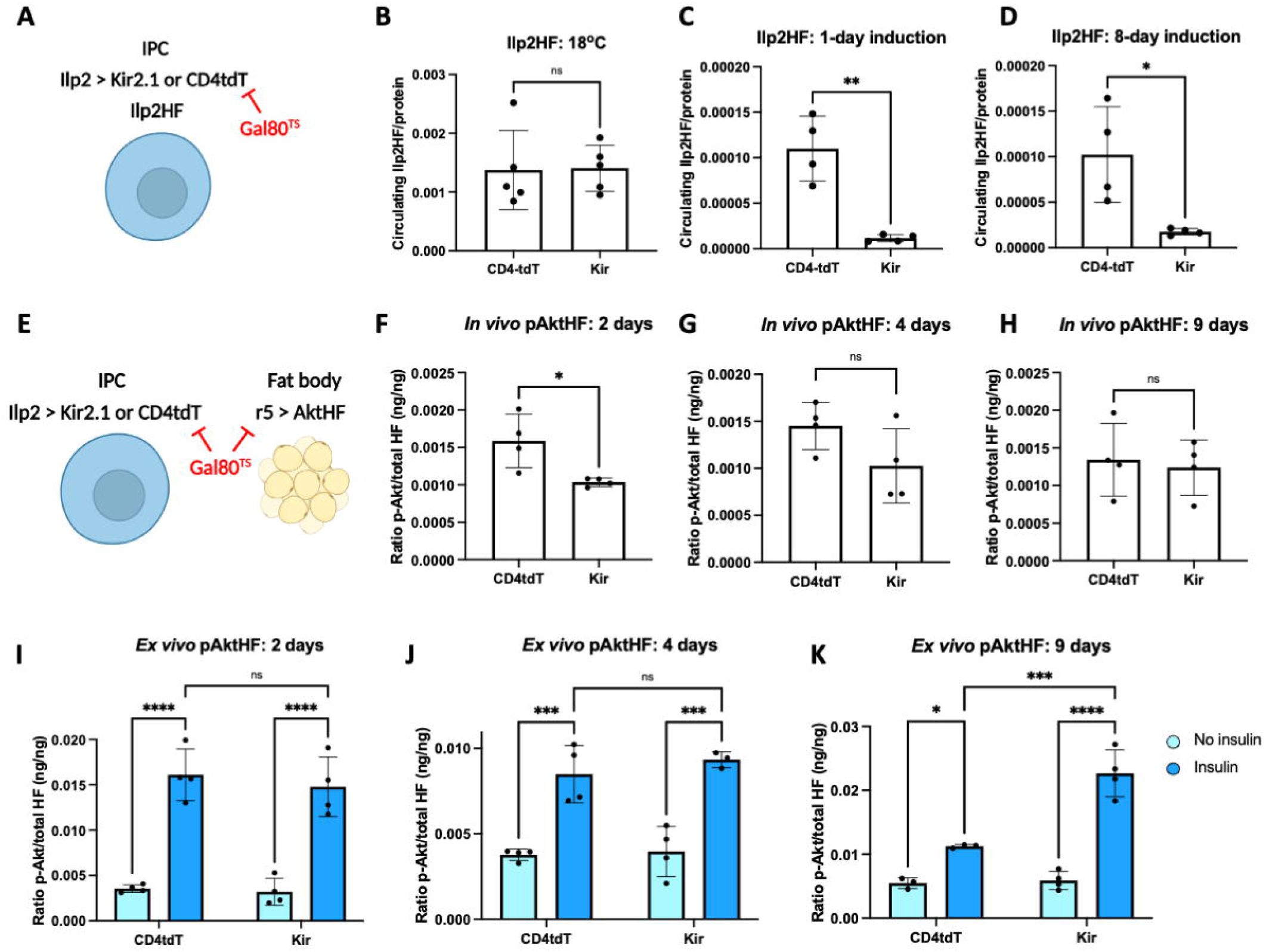
pAktHF ELISA can detect physiological changes of insulin sensitivity. A) Genetics of Ilp2HF measurements (Figure 2B-D), in which insulin-producing cells express either the inhibitory potassium channel Kir2.1 or the control CD4-tdTomato. IPCs also express a knock-in HA- and FLAG-tagged Ilp2 (Ilp2HF) used to measure circulating insulin. Transgene expression of Kir2.1 or CD4-tdTomato is under control of Gal80^TS^. B-D) Measurements of circulating insulin at baseline (18°C, B), following a 1-day induction at 30°C (C), or following an 8-day induction at 30°C (D). E) Genetics of pAktHF measurements (Figures 2F-H), in which IPCs express either Kir2.1 or CD4-tdTomato, while the fat body expresses AktHF. All transgene expression is under control of Gal80^TS^. F-H) Measurements of *in vivo* pAktHF in the adult fat body following 2 (F), 4 (G), and 9 days (H) at 30°C, which permits the expression of Kir2.1 or CD4-tdTomato in IPCs. I-K) *Ex vivo* insulin-stimulated pAkt HF in the adult fat body following 2 (I), 4 (J), and 9 days (K) at 30°C. Human insulin used at 172nM. In all figures, error bars represent the standard deviation. P-values were generated using a two-tailed *t*-test (Figures 3B-D, F-H) or a two-way ANOVA (Figure 3I-K). * indicates *P*<0.05, ** *P*<0.01, *** *P*<0.001 and **** *P*<0.0001. N.S. indicates statistically not significant.

Insulin sensitivity in target organs can adapt to changes of insulin output [1, 3], and our findings suggested that insulin sensitivity in the fat body might be increased in adaptation to chronic hypoinsulinemia. To test this directly, we used the *ex vivo* pAktHF assay to measure fat body responses to insulin in flies with chronic hypoinsulinemia, or in control flies with normoinsulinemia (**Figure 3I-K**). This revealed increased insulin sensitivity in fat body of flies with prolonged IPC silencing and insulin reduction, providing evidence of enhanced insulin sensitivity from chronic hypoinsulinemia. Thus, the combination of these distinct *in vivo* and *ex vivo* assays provides a powerful tool to assess changes in both insulin signaling and sensitivity in metabolic organs over a physiologically-relevant timescale.

### Assessing genetic insulin resistance by measuring AktHF phosphorylation

Our findings suggested the pAktHF ELISA could be used to identify novel genetic regulators of insulin resistance, an established risk for type 2 diabetes [5]. To investigate this possibility, we first assessed pAktHF changes in flies after genetic suppression of known insulin signaling regulators (**Figure 4A**; [12]). After shRNA knockdown in fat body of the insulin receptor (*InR*) or *chico*, the *Drosophila* ortholog of IRS1/2, we observed a significant reduction of pAktHF in *ad libitum* fed flies (**Figure 4B**). As expected, knockdown of *Pten*, a phosphatase inhibitor of insulin signaling, led to increased pAktHF levels, indicating enhanced insulin signaling. By contrast, knockdown of *Rheb*, which encodes a GTP-binding protein downstream of Akt, did not produce significant changes of pAktHF (**Figure 4B**). Since pAktHF dynamics depend on nutrient and insulin (**Figure 1E**), we next assessed AktHF regulation during refeeding of flies with impaired insulin signaling. In flies with fat body loss of *chico* and challenged by fasting and refeeding, we observed blunted AktHF phosphorylation compared to controls (**Figure 4C**); this result supports findings observed in the *ad libitum-*fed state (**Figure 4B**). We found similar changes using our *ex vivo* assay (**Figure 4D**). We also observed similar changes in pAktHF levels after knockdown of *InR, chico, Pten* or *Rheb* in muscle (**Figure 4B, Figure S2)**. These findings provide evidence that the pAktHF ELISA can readily quantify changes in insulin signaling and sensitivity resulting from tissue-specific genetic manipulations.

**Figure 4:**
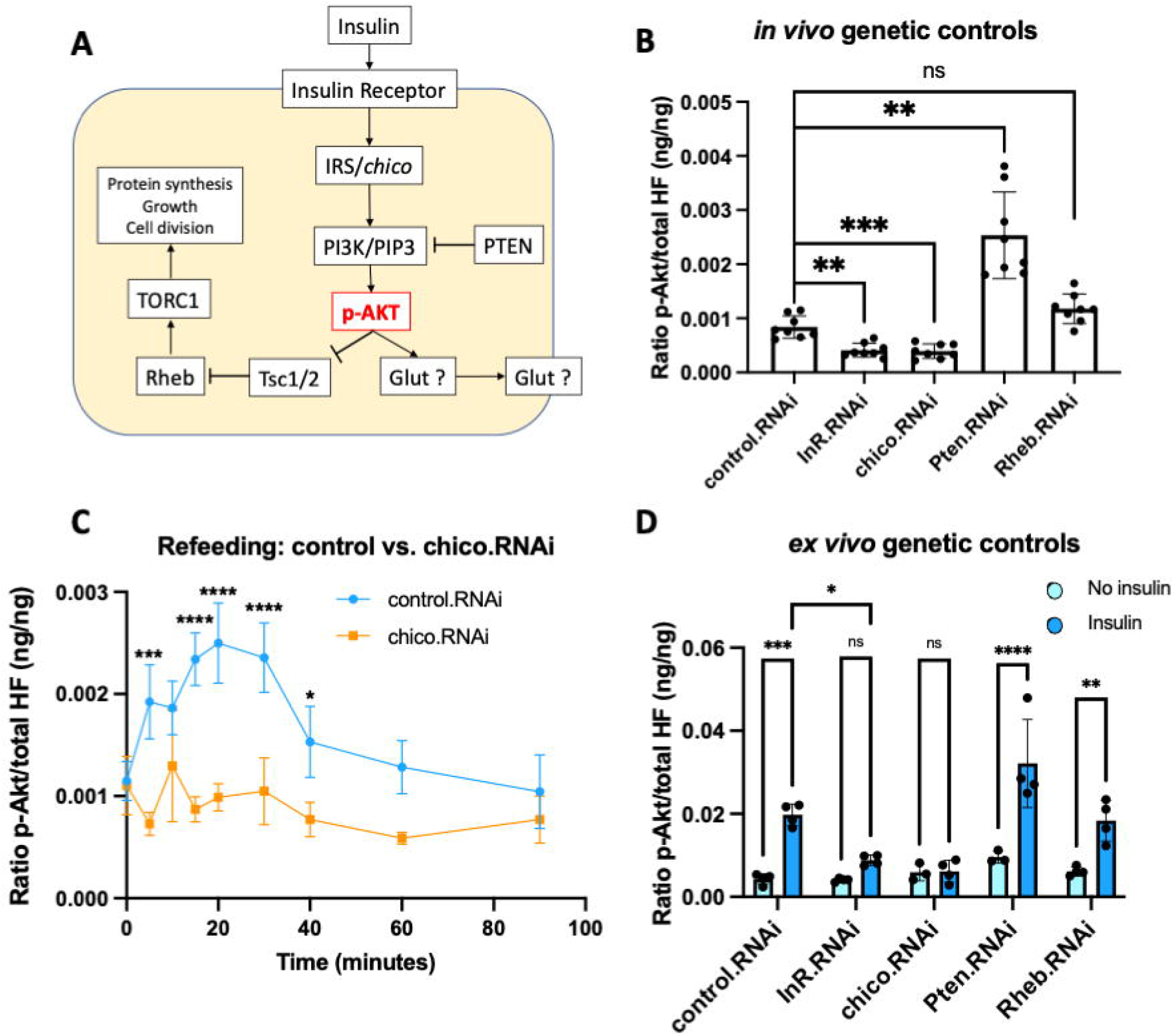
Genetic insulin resistance measured using the pAktHF ELISA A) Schematic of insulin signaling in metazoan cells. B) *in vivo* measurement of pAkt following shRNA knockdown of various known insulin signaling components in the adult fat body (n=8 flies per genotype). C) Oral glucose-stimulated AktHF phosphorylation in control or *chico*.RNAi flies (n=4 flies per time point). D) Induction of pAkt following *ex vivo* insulin stimulation in shRNA knockdown of known insulin signaling components. Human insulin used at 172nM (n=4 fat bodies per bar). P-values were generated using a two-tailed *t*-test (Figures 4B and 4C) or a two-way ANOVA (Figure 4D). * indicates *P*<0.05, ** *P*<0.01, *** *P*<0.001, and **** *P*<0.0001. N.S. indicates statistically not significant.

### Discovering novel regulators of insulin resistance

Our assay provides an opportunity to identify novel regulators of insulin signaling, like *Drosophila* Serine/Threonine (Ser/Thr) phosphatases not previously associated with insulin signaling or Akt regulation. To facilitate efficient genetic screens using publicly available UAS-RNAi lines, we used flies harboring Lpp-Gal4, a fat body specific driver, UAS-AktHF, and Tubulin-Gal80^TS^. The biological activity of UAS-AktHF expressed in the fat body using Lpp-Gal4 was confirmed by measuring pAktHF dynamics during a fasting-refeeding challenge (**Figure S3**), which showed that the rise and clearance of pAktHF resembled that observed using the *LexA/LexAop-AktHF* system (**Figure 1E**). Likewise, UAS-shRNA-mediated knockdown of *InR* or *Pten* produced expected changes in pAktHF compared to *white* or *mCherry* controls (**Figure 5A**).

**Figure 5:**
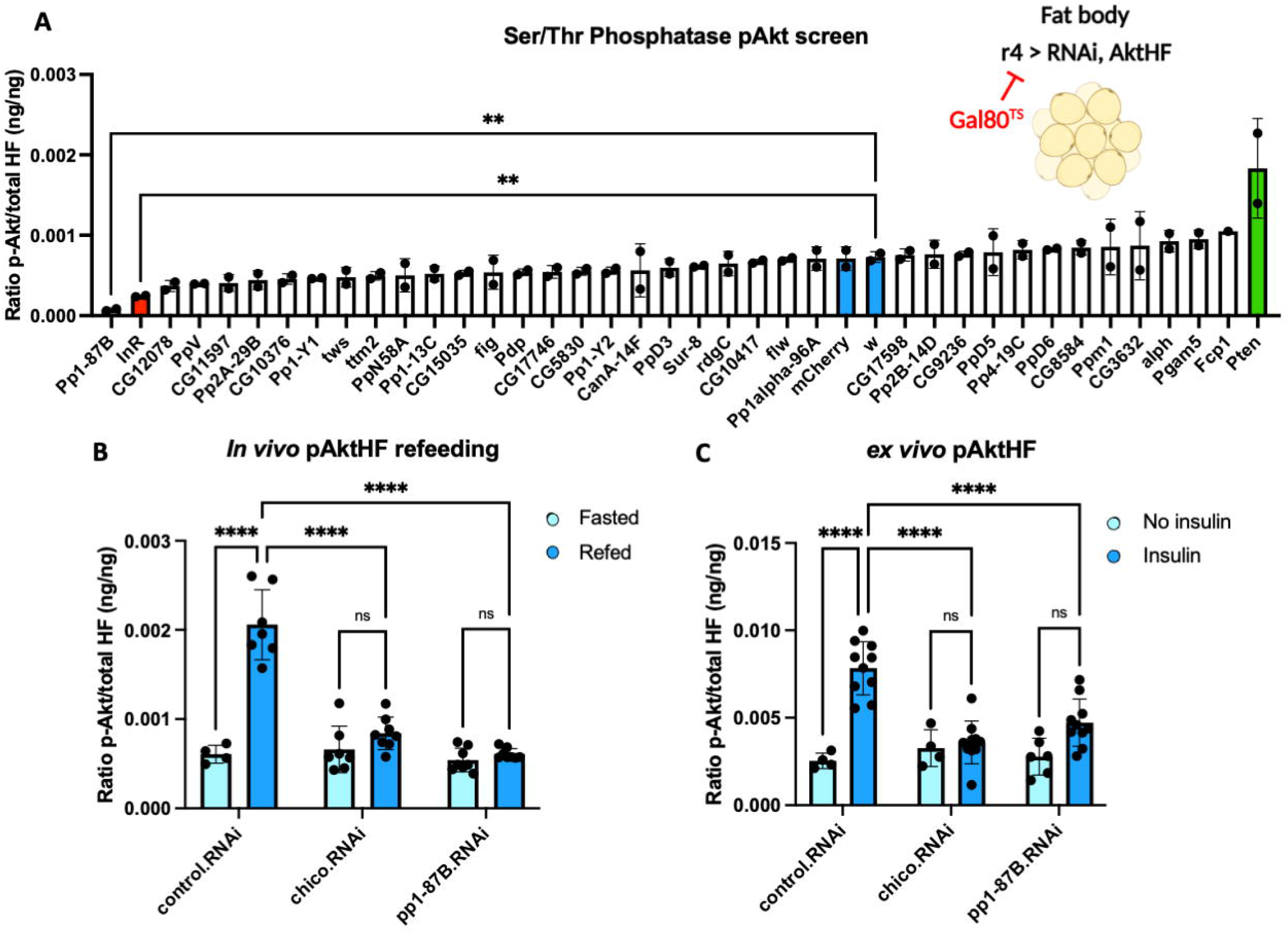
*In vivo* screen of Ser-Thr phosphatases A) *In vivo* pAkt screen of Ser/Thr phosphatase RNAi in the adult fat body. Includes negative controls (mCherry, w.RNAi), upward positive control (Pten.RNAi) and downward positive control (InR.RNAi). n=2 flies per genotype. B) Measurements of *in vivo* pAkt in fasted flies or 20 minutes following an oral glucose challenge. C) Measurements of *ex vivo* pAkt in dissected fat bodies before or after addition of human insulin. P-values were generated using a two-tailed *t*-test (Figure 5A) or a two-way ANOVA (Figures 5B, C). ** indicates *P*<0.01, and **** *P*<0.0001. N.S. indicates statistically not significant.

We then measured the ratio of pAktHF/AktHF after shRNA-mediated knockdown of 35 Ser/Thr phosphatases in *ad libitum* fed flies and found that fat body-specific knockdown of *Pp1-87B*, the putative *Drosophila* ortholog of the alpha and gamma catalytic subunits of human protein phosphatase 1, produced a significant decrease in AktHF phosphorylation (**Figure 5A)**. To validate this *in vivo* finding further, we performed a fasting-refeeding challenge (**Figure S4**). *Pp1-87B* fat body knockdown did not affect fat body pAktHF levels in fasted flies, but these flies had severely impaired pAkt levels following glucose refeeding (**Figure 5B)**. Insulin resistance was measured using the *ex vivo* pAktHF assay, and demonstrated that for a fixed dose of insulin, fat bodies lacking Pp1-87B show significantly diminished responsiveness to insulin stimulation (**Figure 5C**). Together, these results provide index evidence that loss of the *Drosophila* Ser/Thr phosphatase Pp1-87B in the fat body led to impaired insulin signaling.

## Discussion

Work here provides both conceptual and technical advances for studies of insulin signaling. The ELISA assay to measure AktHF or pAktHF quantifies insulin-dependent *Drosophila* Akt phosphorylation on a physiological timescale in single flies. Combination of this assay with powerful binary genetic expression systems additionally permits *tissue-specific* measures of *in vivo* insulin signaling and Akt phosphorylation without dissection; this includes studies of communication between insulin producing cells and insulin target organs, quantification of insulin resistance, and genetic screens to discover regulators of insulin signaling. These findings reveal uses of *Drosophila* to investigate hormonal and physiological regulation relevant to human metabolic diseases like obesity and diabetes.

Here, we demonstrated the insulin dependence of AktHF phosphorylation measured by the pAktHF ELISA, using physiological, genetic and biochemical approaches. Evidence here includes: (1) alignment of the kinetics of circulating insulin and pAktHF excursions in a fasting-refeeding paradigm (**Figure 1E-G)**, (2) demonstration of AktHF phosphorylation in single flies after addition of insulin *ex vivo* (**Figure 2**), and (3) altered pAktHF levels after genetic loss-of-function targeting insulin signal transduction components (**Figure 4**). To the extent that *in vivo* pAktHF measurements in specific tissues reflect endogenous insulin signaling dynamics, we foresee that the sensitivity of the pAktHF ELISA – which permits measures in single flies – should facilitate studies that complement or even substitute for current ELISA measures of circulating fly insulin excursions, which requires pooling of hemolymph from multiple flies [11, 13]. Demonstration here that suppression of insulin secretion from adult IPCs reliably correlated with reduced pAktHF levels in fat body (**Figure 3**) further supports this possibility.

To measure Akt phosphorylation, we labeled *Drosophila* Akt with the immuno-epitopes HA and FLAG, permitting quantification of AktHF or pAktHF, a general experimental strategy we previously used to measure dynamic *in vivo* circulating levels and total levels of Insulin-like peptide 2 [11]. *Drosophila* Akt has multiple regulatory phosphorylation sites [14] and the Ser505 and Thr342 residues have been previously studied [15]. The functional relevance of the vertebrate Akt1-Ser473 residue (which corresponds to *Drosophila* Akt-Ser505) has also been well-studied [16]. Phosphorylation of AktHF-Ser505 was detected with a pSer-specific antibody, a key reagent for building our pAktHF ELISA. Targeting of established ‘upstream’ regulators, like *InR, chico* and *Pten*, validated use of this ELISA to detect and measure changes of insulin signaling strength in adult flies. Prior work suggests that Akt-pSer505 may be regulated by Akt-*dependent* signaling through the Tsc1/Tsc2-TOR-S6K pathway (Kockel et al., 2010), and we have detected pAktHF changes in muscle after shRNA-based suppression of *Rheb*, the GTP binding protein downstream of Akt in the Tsc1/2-TOR pathway (**Figure 4A**). In mammals, Akt-dependent signaling governs phosphorylation of factors like AMPK, and AS160, which regulate glucose transporters like Glut4 [17]; we speculate that development of phospho-protein ELISAs for ERK, AMPK or 4-EBP – analogous to the pAktHF ELISA – could lead to development of additional powerful quantitative assays to measure insulin signaling output at additional signaling detection end points. In addition to phosphorylation, and based on principles identified here and in prior work [11], we speculate that genetic approaches could also generate additional ELISAs to measure other chemically-stable covalent protein modifications governed by signaling relays.

The pAktHF ELISA described here is distinct and complementary to prior assays developed to assess or infer insulin signaling in fruit flies. These include immunohistological staining of phospho-Akt (pAkt) in fixed dissected tissues [9, 10] or imaging of cellular localization of a pleckstrin homology domain-green fluorescent protein [18] to infer insulin receptor-dependent regulation of phosphoinositol-3 kinase (PI3K). While providing valuable insights about insulin signaling regulation, these qualitative or semi-quantitative assays lacked the temporal resolution, tissue specificity, or sensitivity of the pAktHF ELISA. In addition, we show how the pAktHF ELISA can be adapted to permit *ex vivo* exposure to exogenous insulin that uncouples fly insulin signaling from endogenous insulin output, allowing unprecedented evaluation of insulin sensitivity in specific target tissues (**Figures 1-2**).

Here we combined the use of *in vivo* and *ex vivo* pAktHF ELISA to investigate and quantify dynamic changes of insulin signaling sensitivity over a timescale of days resulting from hypoinsulinemia. Prior studies have shown that organisms can compensate for insulin resistance by increasing insulin output, while primary hyperinsulinemia leads to insulin resistance in target tissues [19, 20]. Co-incidence of hypoinsulinemia and insulin hypersensitivity has been noted in fasted flies, Ames dwarf mice, pups of undernourished rats, and offspring of T2DM patients [9, 21-23]. Here, we showed that electrical silencing of insulin-producing cells led to hypoinsulinemia and – initially – to reduced fat body pAktHF levels, as expected. However, pAktHF levels later *normalized* in these flies, suggesting adaptive enhancement of insulin signaling, and we used *ex vivo* pAktHF ELISA to reveal development of fat body hypersensitivity to insulin (**Figure 3**). Together, both assays provide novel measures and evidence of dynamic responses in peripheral insulin target organs to insulinemia.

Studies here also successfully used the pAktHF ELISA to identify genetic regulators of insulin resistance, including established factors (like *InR, chico* and *Pten*) and the phosphatase encoded by *Pp1-87B*. We show that this assay is sensitive to genetic perturbations, whether in single *ad libitum*-fed flies, refeeding-challenged flies, or in response to exogenous insulin. We used the pAktHF ELISA to quantify the ratio of phosphorylated-to-total AktHF, making it well-suited for detecting the impact of post-translational regulators like kinases and phosphatases on insulin signaling. Compared to kinase regulators of insulin signaling [1, 6] phosphatases remain relatively understudied. *Drosophila* genetic screens have revealed phosphatases governing wing disc circadian systems, and embryonic development [12, 24-26]. In a genetic screen of Ser/Thr phosphatases, we found that Pp1-87B suppression potently inhibited Akt phosphorylation. Moreover, this result was confirmed using our *ex vivo* assay, indicating a requirement for Pp1-87B in regulating of insulin-dependent Akt phosphorylation. In contrast to fat body suppression of *Pten*, suppression of *Pp1-87B* led to reduced pAktHF levels, indicating distinct mechanisms might mediate regulation of Akt phosphorylation by Pten and Pp1-87B phosphatases. Pp1-87B is the *Drosophila* ortholog of the alpha and gamma catalytic subunits of human protein phosphatase 1 (PP1), which has been the target of multiple human drug trials (NCT03886662, NCT03027388, NCT01837667) based on its postulated roles in cardiac function, immunity and cancer [27]. PP1 contains a catalytic subunit and one or several additional regulatory subunits which confer tissue and substrate specificity [28, 29]. PP1 is a known *target* of Akt signaling and regulates phosphorylation of glycogen synthase to activate glycogen synthesis [30]. Genetic studies have identified a human T2DM risk variant rs4841132 within the long non-coding RNA *LOC157273* on chr8p23.1, associated with altered expression of *PPP1R3B* which encodes a PP1 regulatory subunit; this suggests a link between PP1 and T2DM or other metabolic traits [31, 32]. However, it remained unknown if PP1 regulates Akt phosphorylation or insulin signaling: genetic suppression of the PP1 regulatory subunit PPP1R3G had no effect on Akt phosphorylation in mouse liver [30]. Findings here motivate future studies of mechanistic links between PP1 and Akt phosphorylation in flies and mammals, and scaled genetic screens to discover additional regulators of insulin signaling.

## Figure Legends

**Figure S1**

A, B) Total levels of AktHF remain constant following refeeding challenge in fat body (A) or muscle (B).

C) Total levels of insulin (Ilp2HF) remain constant following refeeding challenge.

**Figure S2**

*In vivo* measurement of pAkt following shRNA knockdown of various known insulin signaling components in the adult muscle (n=8 flies per genotype). P-values were generated using a two-tailed *t*-test * indicates *P*<0.05, and *** *P*<0.001. N.S. indicates statistically not significant.

**Figure S3**

Oral glucose-stimulated induction of AktHF phosphorylation in adult fat body (Lpp-Gal4) (n=4 flies per time point).

**Figure S4**

Flies of different genotypes immediately following a refeeding challenge with glucose containing blue dye. The ingested food can be visualized in the abdomen. Qualitatively, genotype differences did not affect the amount of food consumed.

## Materials and Methods

### Generation of transgenic lines

To generate p13xLexAop2-IVS-Akt-HF (or LexAop-AktHF), a gene fragment consists of Drosophila Akt protein coding sequence (SD10374) without the stop codon, HA epitope coding sequence, a GGGS flexible hinge coding sequence, and FLAG epitope coding sequence was synthesized (Integrated DNA technologies), and cloned to NotI and XbaI sites on pJFRC19-13xLexAop2-IVS-myr-GFP vector [33]. The construct was inserted to PBac{y^+^-attP-9A}VK00027 site on the third chromosome by integrase-mediated transformation.

To generate pJFRC-MUH-Akt-HF (or UAS-AktHF), the same above gene fragment was cloned to NotI and KpnI sites on pJFRC-MUH vector [33]. The construct was inserted to PBac{y^+^-attP-9A}VK00027 site on the third chromosome by integrase-mediated transformation.

To generate pR5-BPnlsLexA::GADflUw (or r5-LexA), <u>r</u> tetramer [34] was PCR amplified from the genomic DNA of r4-Gal4 line (Bloomington stock #33832) using the primers GCCCTTTCGTCTTCAA<u>GAATTC</u>CTAGTCTTAAAATAATCAGGCGTAGAGTCAGAG and CGGGCGAGCT<u>CGGCCG</u>GCAGTCGACTGATCAGATCTTCGTAGGCC, and the largest PCR fragment was cloned to EcoRI and NaeI site on pBPnlsLexA::GADflUw vector [33]. Upon sequencing the construct, we found five repeats of 80 bp *Yp1* <u>r</u> enhancer were cloned, and thus named the construct as ‘r5-LexA’. The construct was inserted to P{CaryP}attP40 site on the second chromosome by integrase-mediated transformation. r5-LexA expression was confirmed by crossing to 13xLexAop-tdTomato.nls (Bloomington stock #66680) with minimal expression in adult optic lobes and restricted expression in abdomen and head fat body tissues, similar to r4-Gal4.

To generate pMhc.F3-580-BPnlsLexA::GADflUw (or Mhc-LexA), the adult leg muscle specific Mhc.F3-580 enhancer [35] was PCR amplified from the genomic DNA of y^1^ w^1118^ control flies using the primers ATGTTACAATT<u>GAACTT</u>ATACCAACTCATTGGCTTTACAAG and CAGAAT<u>CTCGAG</u>AGTCTCCCCTCTTACAACGATGTC, and cloned to EcoRI and XhoI (NaeI site modified) sites on pBPnlsLexA::GADflUw vector [33]. The construct was inserted to P{CaryP}attP40 site on the second chromosome by integrase-mediated transformation.

To generate pDilp215-1-BPnlsLexA::GADflUw (or Ilp2-LexA), 541 bp Dilp215-1 enhancer [36] from pDilp215-1-H-Stinger [37] was cloned to EcoRI and XhoI(NaeI site modified) sites on pBPnlsLexA::GADflUw vector [33]. The construct was inserted to P{CaryP}attP40 site on the second chromosome by integrase-mediated transformation.

To generate pDilp215-1-BPGUw (or Ilp2-Gal4), 715 bp EcoRI – KpnI fragment containing 541 bp Dilp215-1 enhancer and 155 bp Drosophila synthetic core promoter from pDilp215-1-BPnlsLexA::GADflUw was cloned to EcoRI and KpnI site on pBPGUw vector [33]. The construct was inserted to P{CaryP}attP2 site on the third chromosome by integrase-mediated transformation.

To generate Ilp2^HF^ knock-in allele (or Ilp2HF) in Ilp2 locus, two 20 bp guide RNA sequences, AGGCGAACTCGCCAACGGCA and TTACGCATGGCGCGCTTGTG, were cloned to pCFD3 vector (Addgene #49410). The donor construct pattB-Ilp2HF-DsRed-SCL was generated by inserting PBac{ScarlessHD-DsRed}module (Addgene #64703) into the TTAA sequence within the C-chain region of pBDP2-gd2HF [10]. The donor and two gRNA constructs were co-injected to vas-Cas9 embryos (Bloomington stock #55821), and F1 progeny were screened for DsRed eye color to identify successful knock-in events. PBac{ScarlessHD-DsRed} module was removed by Tub-PBac transposase (Bloomington stock #8285). The resulting Ilp2HF allele was sequenced to verify the reversion of TTAA sequence within the C-chain region. We note that Ilp2HF allele does not contain the endogenous Ilp2 intron, and Ilp2HF transcript level is about six times higher than wild type Ilp2 expression overall in 1, 5, and 10 day old males determined by qPCR.

### Drosophila strains and husbandry

For LexA phospho-Akt experiments flies were generated on a *yellow white* (yw) background, with the following genotype: yw ; r5-LexA, Tubp-Gal80TS ; Ilp2-Gal4, LexAop-Akt-HF. These flies were crossed to LexAop-RNAi lines [12]. For Gal4 phospho-Akt experiments, flies were generated on a *yellow white* (yw) background, with the following genotype: yw ; Ilp2-LexA, Tubp-Gal80TS ; Lpp-Gal4, UAS-Akt-HF. These flies were crossed to UAS-RNAi lines obtained from the Bloomington *Drosophila* Stock Center. For Ilp2HF refeeding experiments, yw ; + ; Ilp2HF, Ilp2-Gal4 flies were used.

Virgin female flies were used for all experiments. All flies were raised on a standard molasses and yeast-containing food media.

### Fasting and refeeding experiments

Flies were fasted for 28-70 hours in vials containing 2% agar. Fasting times depended on the age and genotype of the fly but were generally fasted until the first flies began to die of starvation to ensure robust refeeding. Flies were refed for 5 minutes on 250mM glucose, 2% agar, 1% blue food colouring. Following refeeding, they were returned to fasting vials. To collect flies for the phospho-Akt refeeding time course, 4 flies were shaken from the vials onto a CO_2_ pad, such that the rest of the flies in the vial remained awake. To reduce handling and stress to the animals, flies were returned to two separate fasting vials which were shaken on alternate time points. Robust refeeding was verified by the presence of blue colour in the abdomen. Single flies were placed in individual 1.5mL Eppendorf tubes and flash frozen in a solution of dry ice and 90% ethanol, before being transferred to -80°C until the conclusion of the time course. To collect flies for the Ilp2HF refeeding time course, flies were separated into separate vials for each time point to avoid repeated anesthetization.

### Ex vivo insulin stimulation

Fat bodies were isolated from live flies anesthetized on CO_2_ by isolating the abdomen and removing the ovaries and digestive tract. Dissections were performed in a drop of PBS on a glass slide. The fat body was placed in a 1.5mL Eppendorf tube with 100µl PBS. To stimulate insulin signaling, human insulin (Sigma-Aldrich I9278) was added at 172nM, unless otherwise indicated, and the tube was mixed by flicking. Unless otherwise specified, fat bodies were incubated with or without insulin for 10 minutes at 22°C.

### p-Akt ELISA

We coated wells in Nunc-Immuno modules (Thermo Scientific 468667) with 100µl of anti-FLAG antibody (Sigma-Aldrich F1804) diluted in 0.2 M sodium carbonate/bicarbonate buffer (pH9.4) to a final concentration of 2.5 µg/ml, then incubated for 16 hours at 4°C. The plate was then blocked with 350 µl of PBS containing 4% bovine serum albumin (Fisher Scientific BP1600) for 5 minutes at 22°C. The plate was washed three times with PBS containing 0.1% Triton-X.

A single intact fly or a single dissected fat body was placed on ice in 100 µl PBS with 1% Triton-X-100, 100mM sodium fluoride, and 5mM sodium orthovanadate. A pestle and cordless motor were used to lyse the tissue. The samples were centrifuged at 21,000g for 30 seconds at 4°C, and the supernatant was used for the ELISA.

For phosphorylated Akt measurement, supernatant was mixed with 50µl of Superblock containing 0.1% Triton X-100 and Phospho-Akt (Ser473) (D9E) XP Rabbit mAb (Cell Signaling Technology #4060) at a 1:2000 dilution. For total Akt measurement, supernatant was mixed with 50µl of Superblock containing 0.1% Triton X-100 and anti-HA-Peroxidase 3F10 antibody (Roche 12013819001) at a 1:10,000 dilution. Sample volumes for specific experiments are tabulated in Supplemental Methods. An equal volume of phospho-Akt standard peptide (PLFPQFpSYQGDYKDDDDK, 0 to 0.5ng/mL) or FLAG(GS)HA standard peptide (DYKDDDDKGGGGSYPYDVPDYA, 0 to 2.5ng/mL) was also loaded onto separate wells. The wells were sealed with an adhesive film (Bio-Rad MSB1001) and incubated on a rotary shaker at 4°C overnight.

Phospho-Akt wells were washed 6 times with PBS containing 0.1% Triton X-100. 100uL of PBS with 0.1% Triton X-100 and 4% BSA containing donkey anti-Rabbit-HRP (Thermo Fisher Scientific A16035) diluted 1:4000 was added to each well and incubated at room temperature on a rotary shaker for 1 hour. Phospho- and total-Akt wells were then all washed 6 times with PBS containing 0.1% Triton X-100. 100 µl of 1-Step Ultra TMB ELISA Substrate (Thermo Scientific 34029) was added to each well and incubated for 15 minutes at room temperature. The reaction was stopped by adding 50µl of 2M sulfuric acid, and the absorbance at 450 nm was immediately measured on a Varioskan™ LUX multimode microplate reader (Thermo Scientific VL0000D0).

### Ilp2HF ELISA

Circulating insulin was measured as previously described [11, 13], with some small modifications. For flies homozygous for Ilp2HF, 6 flies were pooled in 57µl of extraction buffer (PBS with 0.1% Triton X-100). For flies heterozygous for Ilp2HF, 8 flies were pooled instead. Hemolymph was extracted on ice for 30 minutes. 42µl of extraction buffer was loaded for circulating Ilp2HF. Flies were then homogenized in 600uL (homozygous for Ilp2HF) or 800uL (heterozygous for Ilp2HF) extraction buffer, and 5µl supernatant was loaded for total Ilp2HF.

## Supplementary methods

### Dilutions and loading volumes for different drivers

Different tissue-specific drivers led to different levels of AktHF expression, so the volumes and dilutions of samples were adjusted to remain in the linear detection range of the assay (Figure 1C).

| Driver |  | Loading volume, total<br>AktHF (μL) | Loading volume,<br>pAktHF (μL) |
| --- | --- | --- | --- |
| r5-LexA | In vivo | Diluted 1:200, loaded 10 | 20 |
|  | Ex vivo | Diluted 1:40, loaded 10 | 20 |
| Lpp-Gal4 | In vivo | Diluted 1:100, loaded 2.5 | 5 |
|  | Ex vivo | Diluted 1:50, loaded 2.5 | 5 |
| Mhc-LexA | In vivo | Diluted 1:100, loaded 20 | 20 |

## Acknowledgments

We thank the Bloomington Drosophila Stock Center (NIH P40OD018537) and the TRiP at Harvard Medical School (NIH/NIGMS R01-GM084947) for transgenic fly stocks used in this study. We thank Ms. E. Wesel for assistance in statistical analysis, and members of the Kim group for advice and encouragement. D.D.T. is a trainee in the Berg Scholars Program at Stanford University School of Medicine, and K.R.C. was supported by a Stanford Vice Provost Undergraduate Education award. Work in the Kim laboratory was supported by NIH awards (R01 DK107507; R01 DK108817; U01 DK123743; P30 DK116074 to S.K.K), the Reid Family, Sadie and Kelly Skeff, H.L. Snyder Foundation and Elser Trust, the Mulberry Essence Foundation, two anonymous donors, and the Stanford Diabetes Research Center (SDRC).

## Author Contributions

DDT, SP and SK designed experiments, DDT, KRC and SP collected data, analyzed data, and DDT, SP and SK wrote the manuscript; LK provided materials and advice; SP and SKK conceived the project; SKK supervised.

