## Supplementary figures and images for "A genetic strategy to measure insulin signaling regulation and physiology in *Drosophila*"

### Supplemental Figure 1

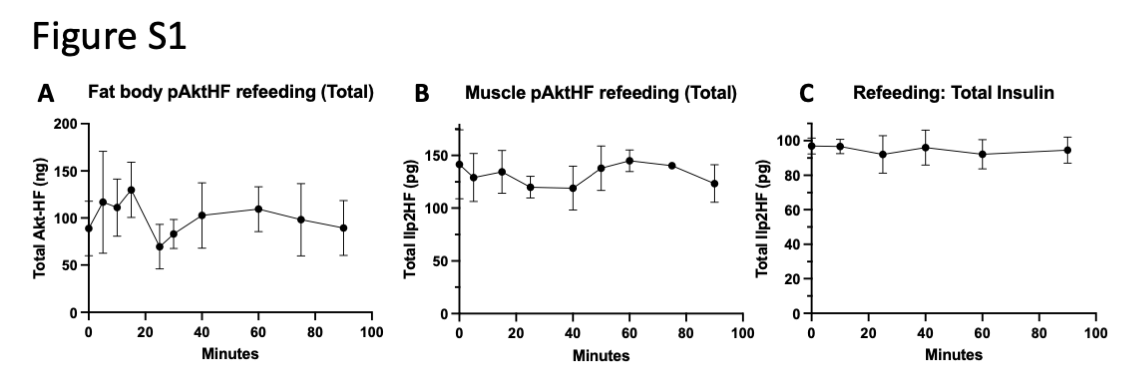

### Supplemental Figure 2

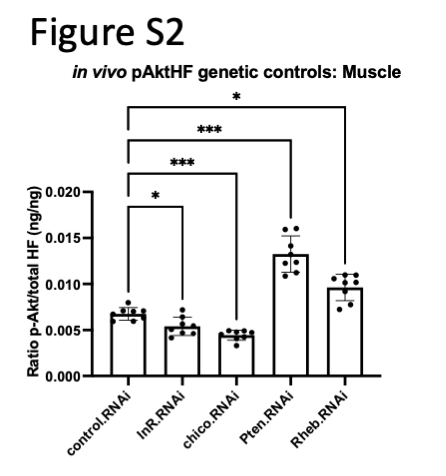

### Supplemental Figure 3

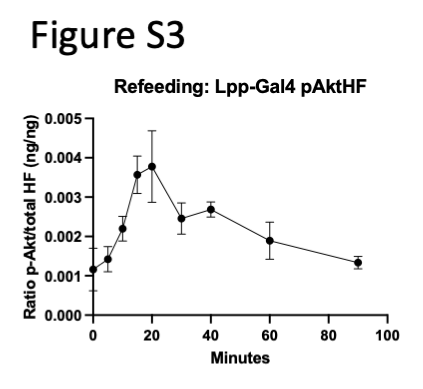

### Supplemental Figure 4

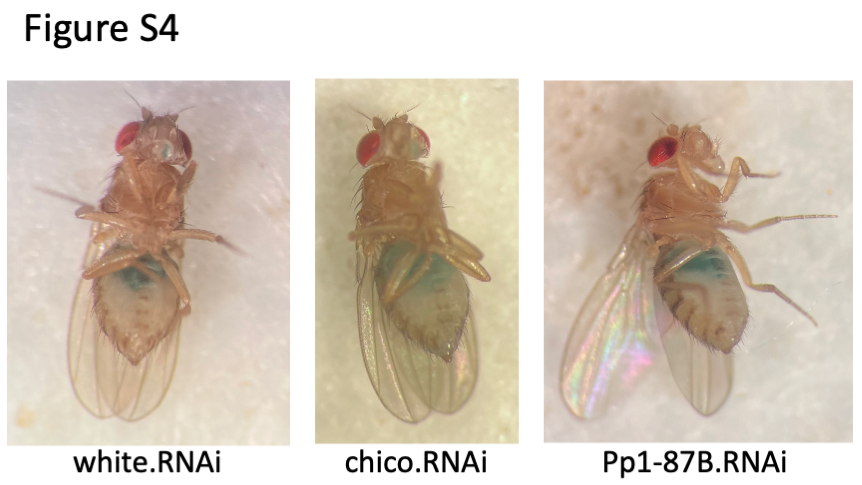
